# Green Synthesis of Different Zein nanoparticles – Characterization and *In vitro* Release studies

**DOI:** 10.1101/2025.09.01.673414

**Authors:** A. T. Dhanya, N. Vaagdevi, M. Archana, Santosh Kanade

## Abstract

Nanoparticle-based drug delivery has gained traction in the biomedical field, as it reduces drug susceptibility, improves bioavailability, and enhances therapeutic efficacy and drug targeting. Among the various types of nanoparticles, biopolymeric nanoparticles composed of proteins and polysaccharides offer a significant advantage in drug delivery due to their biocompatibility, biodegradability, environmental sustainability, and versatility. Nanoparticles made of zein, a prolamine class of protein, are widely studied for their use in the biopharmaceutical industry. However, the clinical translation of the drug-loaded zein nanoparticle remains a challenge. Different zein nanoparticles were prepared using the green synthetic approach by incorporating different polysaccharides, sodium alginate, and pectin to improve their characteristics and drug-release properties. Herbal extract was chosen as the drug material for the drug release study. Standard methods calculated the nanoparticle yield, percentage of encapsulation, and drug content. The surface morphology and size of the nanoparticles were studied by Scanning Electron Microscope (SEM); structural characterization by Fourier Transform Infra-Red and UV-visible spectroscopic methods, and thermal stability by Differential Scanning Calorimetry (DSC) and Thermogravimetric analysis (TGA). The *in vitro* drug release was performed using the dialysis method. The SEM results specify the formation of spherical nanoparticles of the appropriate size. The spectroscopic studies confirmed the incorporation of all the components in the nanoparticles. DSC and TGA analysis confirmed the thermal stability of the nanoparticles. The drug release studies demonstrated diffusion-based drug release, which followed first-order kinetics and the Higuchi mechanism. From the results, it was confirmed that the incorporation of polysaccharides improves the encapsulation efficiency and the drug-release properties of the nanoparticles. It was also evident that the encapsulation of the herbal extract further improved the stability of the nanoparticles.

**Highlights:**

- Zein nanoparticles were prepared from natural sources using a green synthetic approach
- Spherical nanoparticles with the appropriate size were formed
- The nanoparticles exhibited good thermal stability
- All the nanoparticles obeyed first-order release kinetics
- According to Higuchi kinetics, the drug release of ZAEA np>ZPEA np>ZEA np, which is in agreement with the Korsmeyer Peppas kinetics model, indicates the value of ‘n’, the release exponent, ZEA np = -0.042; ZAEA np = 0.49; ZPEA np = 0.27
- ZAEAnp exhibited better encapsulation and release properties

## Introduction

Nanoparticles are the magic bullets in drug delivery as they improve targeting efficiency, bioavailability, biodistribution, therapeutic index, pharmacokinetics, controlled drug release, and reduced toxicity due to their size, shape, surface features, and drug loading [1]. To date, different types of synthetic and biodegradable polymers, metals, silica, etc, and liposomes have been used for this purpose [2–5]. However, biopolymeric nanoparticles have captured significant interest in drug delivery applications due to their biocompatibility, biodegradability, stability, and non-toxic characteristics[1]. Several proteins and polysaccharides, like albumin, zein, cellulose, starch, pectin, sodium alginate, etc, serve the purpose of nanoparticle synthesis[6–12]. Protein and polysaccharide molecules, in suitable conditions, assemble into particles through electrostatic attraction between oppositely charged groups. Protein has a specific property called isoelectric point (pI), which is the pH at which it possesses no charge but has localized regions that can significantly influence its interactions with ionic polysaccharides. Above their PI proteins exhibit a net negative charge, and a net positive charge below it. The interactions between the anionic polysaccharides and cationic protein surfaces or vice versa can lead to the formation of protein-polysaccharide complexes. This property can be utilized for the green synthesis of biopolymer structures like nanoparticles for various applications, like biomedicine[13,14].

Green synthetic methods have gained momentum, and plant metabolites and sources are utilized in the synthesis of nanoparticles for various applications[15]. As nanoparticles are widely used in biomedical applications, there is more interest in developing processes that use nontoxic, inexpensive chemicals, low-energy-consuming, and environment-friendly methods for their synthesis, and green synthesis becomes an alternative to chemical or physical methods[16]. Nanoparticles are synthesized using a one-step procedure in the green synthetic method and were found to have diverse characteristics, higher stability, and appropriate dimensions[17].

Among the various protein nanoparticles, developing zein-based nanoparticles has become a research hotspot as they enhance the pharmaceutical properties of drugs and exhibit low immunogenicity [18]. Zein is a prolamine protein class found in corn and maize. It is included in the generally recognized as safe (GRAS) category of proteins by the United States Food and Drug Administration [19]. Its hydrophobicity, alcohol solubility, and biocompatibility make it an important candidate for developing nanoparticles for the drug delivery of hydrophobic drugs. Due to their self-assembly nature, these particles can be prepared effortlessly using zein [19–21]. Zein nanoparticles are known to be unstable and prone to aggregation, primarily because of their hydrophobic properties and charge characteristics [22]. These limitations can be addressed by utilizing stabilizing agents like polysaccharides. Polysaccharides can function as emulsifiers and stabilizers, forming a protective coating around the zein nanoparticles and thereby enhancing their dispersibility and stability [22,23].

One of the polysaccharides used for modifying zein nanoparticles was sodium alginate, a natural anionic polysaccharide found in seaweed or brown algae. Sodium alginate is a linear copolymer of the homopolymers, (1 4) β-D-mannuronopyranosyl and (1 4) α-L-guluronopyranosyl units [18–20]. Another polysaccharide used was pectin, a cell wall polysaccharide found in plants. Pectin contains polygalacturonic acid units with methyl-esterified carbonyl groups. The degree of esterification (DE) varies in pectin from different sources, and accordingly, they are classified into high methoxyl pectin (HM) with DE higher than 50% and low methoxyl pectin (LM) with DE less than 50% [11,21]. Several studies indicate the importance of sodium alginate and pectin in stabilizing the other components of the drug delivery system, such as zein, chitosan, inulin, etc[11,22–24]. Due to their structure with egg box regions and charge difference, sodium alginate and pectin can undergo electrostatic interactions or crosslinking with other biomaterials to form complexes [18,19,21,25]. In the presence of an emulsifying agent and surfactant, silicone oil and these components complex to form nanoparticles [26,27]. In this context, our research focuses on the green synthesis of different types of zein nanoparticles using biopolymers zein, sodium alginate, and pectin from natural sources and studies on their drug-release properties.

## Materials and Methods

### Chemicals and reagents

Pectin (Cat No. 9000-69-5) and Silicon oil (Cat No. 63148-62-9) were purchased from Sisco Research Laboratories, Zein (Cat No. 9010-66-6) from Hi Media Chemicals, and Alginate (Cat No. 9005-38-3) from Duchefa Biochimie. All reagents used were of analytical grade.

### Plant Collection

*Wrightia tinctoria* plant leaves were collected from Kuttamassery, Ernakulam, Kerala, India. Prof. Dominic, St. Teresa College, Kochi, Kerala, India, authenticated the collection. The Herbarium was submitted to the Department of Plant Sciences, University of Hyderabad, with accession number UH1002.

### Preparation of Plant Extract

**Leaves** were collected, air dried, and powder was subjected to Soxhlet extraction with 80% methanol. The methanolic fraction was then fractionated with petroleum ether, diethyl ether, and ethyl acetate [11,28,29]. The ethyl acetate (EA) fraction was selected for the green synthesis of biopolymeric nanoparticles and is referred to as EA.

### Green synthesis of biopolymeric nanoparticles

Biopolymeric nanoparticles were prepared by ultrasonication using varying concentrations of zein, pectin, alginate, and ethyl acetate extract of *Wrightia tinctoria* leaves [11].

### Synthesis of Zein (Z - np) and EA-Zein Nanoparticles (ZEA-np)

Ten milliliters of 0.01% silicone oil was sonicated at 20% amplitude, followed by the abrupt addition of 2 milliliters of zein (0.5 mg/mL) in 80% ethanol to the silicone oil emulsion. Sonication was continued for another 15 minutes. ZEA-nps were prepared as reported in [11] with some modifications. One milliliter of zein (0.5 mg/mL) and 1 milliliter of extract (5 mg/mL) were mixed and added abruptly to the silicone oil emulsion, and then sonicated for 15 minutes at 20% amplitude.

### Synthesis of Zein - Alginate (ZA - np) and EA-Zein - Alginate Nanoparticles (ZAEA - np)

Ten ml of 0.01% silicone oil was sonicated at 20 % amplitude with the addition of 2 ml of zein (0.5mg/ml) in 80% ethanol to the emulsion. After 15 minutes, 2 ml of Alginate solution (0.1%) was added dropwise, and sonication was continued for 25 minutes. ZAEA-np’s were prepared by adding the mixture of 1 ml of zein (concentration) and 1 ml of extract to the silicone oil emulsion with sonication for 15 minutes at 20% amplitude, followed by the addition of 2 ml of 0.1% Alginate solution dropwise with sonication for another 25 minutes.

### Synthesis of Zein - Pectin (ZP - np) and EA-Zein-Pectin Nanoparticles (ZPEA -np)

Ten ml of 0.01% silicone oil was sonicated at 20 % amplitude, and 2 ml of zein (0.5mg/ml) in 80% ethanol was added to the silicone oil emulsion. After 15 minutes of sonication, 2 mL of 0.05% pectin solution was added dropwise, and sonication was continued for another 25 minutes. ZPEA-np’s were prepared by a mixture of 1 ml of zein (0.5mg/ml) and 1 ml of extract (5mg/ml) added abruptly to the silicone oil emulsion and sonicated for 15 minutes at 20% amplitude, followed by the addition of 2 ml of 0.05% pectin solution dropwise, and sonication was continued for another 25 minutes.

### Nanoparticle recovery (%), Encapsulation efficiency (%) and Drug content (%)

The percentage of recovery, encapsulation efficiency, and drug content of the drug-encapsulated nanoparticles were calculated by standard methods [11,30]. The percentage of nanoparticle recovery was calculated using the following equation (1)

### Nanoparticle recovery (%)

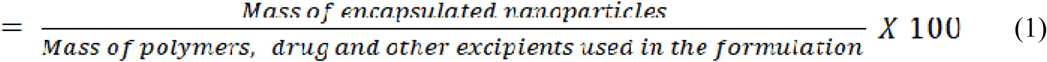

The mass of encapsulated nanoparticles is the weight of nanoparticles formed, and the mass of polymeric nanoparticles is the weight of the particles prepared without drug.

The encapsulation efficiency (%) and drug content (%) were calculated using equations (2) & 3

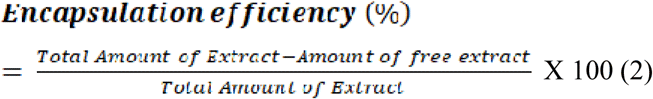

The total amount of extract is the amount used in nanoparticle preparation; the amount of free extract is calculated by analysing the absorbance of the supernatant after sedimentation of nanoparticles by centrifugation. The absorbance of the supernatant was taken at 388nm, and the amount of the free extract is calculated from the calibration curve of the extract at 388nm.

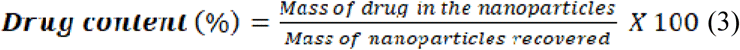

### Characterization of Nanoparticles

The nanoparticles were characterized for their surface morphology and size with Scanning Electron Microscopy (SEM). Particle size and zetapotential measurements were analysed with the Dynamic Light Scattering (DLS) method. Spectroscopic methods, such as UV-Visible and FT-IR spectroscopy, were employed to confirm the incorporation of all components in the nanoparticles. Thermal stability was analysed using Thermo Gravimetric Analysis (TGA) and Differential Scanning Calorimetry (DSC) methods.

### Sem Analysis

The surface morphology, size, and shape of the nanoparticles were analyzed with a Scanning Electron Microscope (SEM EDAX) (CARL ZEISS Ultra 55; EDX: Oxford, INCAx-act, Germany). The particles were coated on carbon tape and were splutter-coated with gold particles for 30 minutes. The sample was then visualized under FE-SEM at a 200nm scale.

### Dynamic Light Scattering

The particle size and zeta potential measurements were conducted using a Zetasizer Anton Paar instrument.

### UV-Visible Spectroscopy

UV–Visible spectroscopy (Jasco V-750 spectrophotometer) was used to compare the absorption spectra of individual components, including zein, sodium alginate, pectin, EA extract fraction, and the nanoparticles ZEAnp, ZAEAnp, and ZPEAnp. A spectral scan was conducted in the 200 – 700nm range. The components, Zein and EA extract, were solubilized in 80% ethanol. Sodium alginate and pectin were solubilised in water. The spectral scan was performed in the respective solvents following baseline calibration using the same solvents.

Among the nanoparticles, ZEAnp was solubilised in 80% ethanol for the spectral scan, as all the components were soluble only in 80% ethanol. As the other two nanoparticles, ZAEAnp and ZPEAnp, contain components that are insoluble in 80% ethanol, the spectral scan was conducted in both water-solubilised and 80% ethanol-solubilised nanoparticles. Before the spectral scan, the nanoparticles were vortexed in respective solvents and centrifuged to separate insolubilised particles.

### FT-IR Spectroscopy

The FT-IR spectroscopic method (Thermo Fischer Scientific, Nicolet iS5 ATIR spectrophotometer) was utilized to analyze the nanoparticle component’s incorporation. The dried samples were used for the analysis.

The thermal stability of the nanoparticles was confirmed with thermogravimetric analyses and differential scanning calorimetric studies.

### Evaluation of *in vitro* drug release and kinetics

Release properties of the extract-incorporated nanoparticles were studied by the dialysis method using a Dialysis membrane with a molecular cut-off of 12,000 KD (Dialysis membrane – 70, HiMedia) [11,30,31]. The dialysis membrane was activated before the experiment. 2M phosphate buffer at pH 7.4 was used. Approximately 500 mg of nanoparticles were suspended in 2 mL of phosphate buffer was added to the dialysis bag. It was dipped in 100 mL of phosphate buffer in beakers. After every hour, 1 ml of buffer was withdrawn and replaced with fresh buffer. OD of the dialysate was measured at 388nm. The cumulative percentage of drug release was calculated and plotted against time in hours to understand the release pattern. The release kinetics was fitted in kinetic models such as zero order, first order, second order, Higuchi model, Korsmeyer Peppas model, and Weibull model using R programming [32].

Statistical analysis:

## Results and Discussion

### Preparation and characterization of nanoparticles

The nanoparticles were prepared using the protein zein from corn and polysaccharides such as alginate and pectin. The side chains of zeins are rich in hydroxyl, carbonyl, and amino groups. Zein is a biocompatible protein that can self-assemble to form nanospheres upon optimizing the solvent polarity. Alginate and pectin have hydroxyl and carboxyl groups as side chains. The nanoparticle is prepared by the ultrasonication method. Ultrasonication is one of the most suitable green ways for nanoparticle synthesis as it saves energy and speeds up the reaction and reactivity[32]. It can also degrade the long polymeric chains of zein, pectin, and alginate most economically and simply. The degradation of the polymers can also be controlled by controlling various factors such as temperature, polymer concentration, frequency, intensity, and time of sonication [33]. Ultrasonication, along with surfactants, facilitates the formation of nanoparticles. In this study, silicone oil or siloxanes are used as surfactants and emulsifying agents. Silicone oil exhibits several characteristics that make it an excellent choice in nanoparticle synthesis. Silicone oil is thermodynamically stable, highly lubricant, has lower surface tension, and is odorless and non-toxic [34]. The yield, nanoparticle recovery percentage, EE percentage, and drug content of nanoparticles were calculated using standard methods [11,30]. The results are given in Table 1. Results indicate that ZAEAnp shows the highest encapsulation efficiency and drug content percentage compared to ZEAnp and ZPEAnp.

**Table 1:** Nanoparticle parameters.

|  | **ZEAnp** | **ZAEAnp** | **ZPEAnp** |
| --- | --- | --- | --- |
| **Nanoparticle yield** | 45.5 ± 0.12mg | 7mg | 55.2 ± 0.16mg |
| **Nanoparticle recovery %,** | 77.4 ± 0.25% | 11.7% | 92 ± 0.27% |
| **% Encapsulation efficiency** | 6.3 ± 0.4% | 32.24 ± 0.23% | 20.46 ± 0.016% |
| **% Drug content** | 0.9 ± 0.06 % | 23 ± 0.16% | 1.85 ± 0.004% |

### Characterization of nanoparticles

Prepared nanoparticles were characterized for shape and surface morphology, particle size, zetapotential, drug entrapment efficiency, and polymer–polymer interaction. SEM characterized the shape and surface morphology of the carrier system. Drug release efficiency was estimated by the equilibrium dialysis method.

### SEM Analysis

The SEM images representing the difference in the shape, size, and surface morphology of different nanoparticles are presented in Fig.1. The size range of different nanoparticles is shown in Table No. 2, which indicates the approximate size range (approximately 100 – 1000 nm). SEM analysis showed that all the nanoparticles were formed in a spherical shape with a smooth surface. Znp and ZEAnp formed better nanoparticles with an appropriate size (Znp 93 – 360 nm and ZEAnp 44 – 256 nm) and less agglomeration. Among the core-shell nanoparticles, ZAnp and ZPnp, ZAnp formed particles with less agglomeration. Still, on extract incorporation, ZAEAnp exhibited aggregation, and the size was also increased from 48 – 400 nm to 400 – 1007 nm. On comparing ZAEAnp and ZPEAnp, ZPEAnp shows better size.

**Figure 1:**
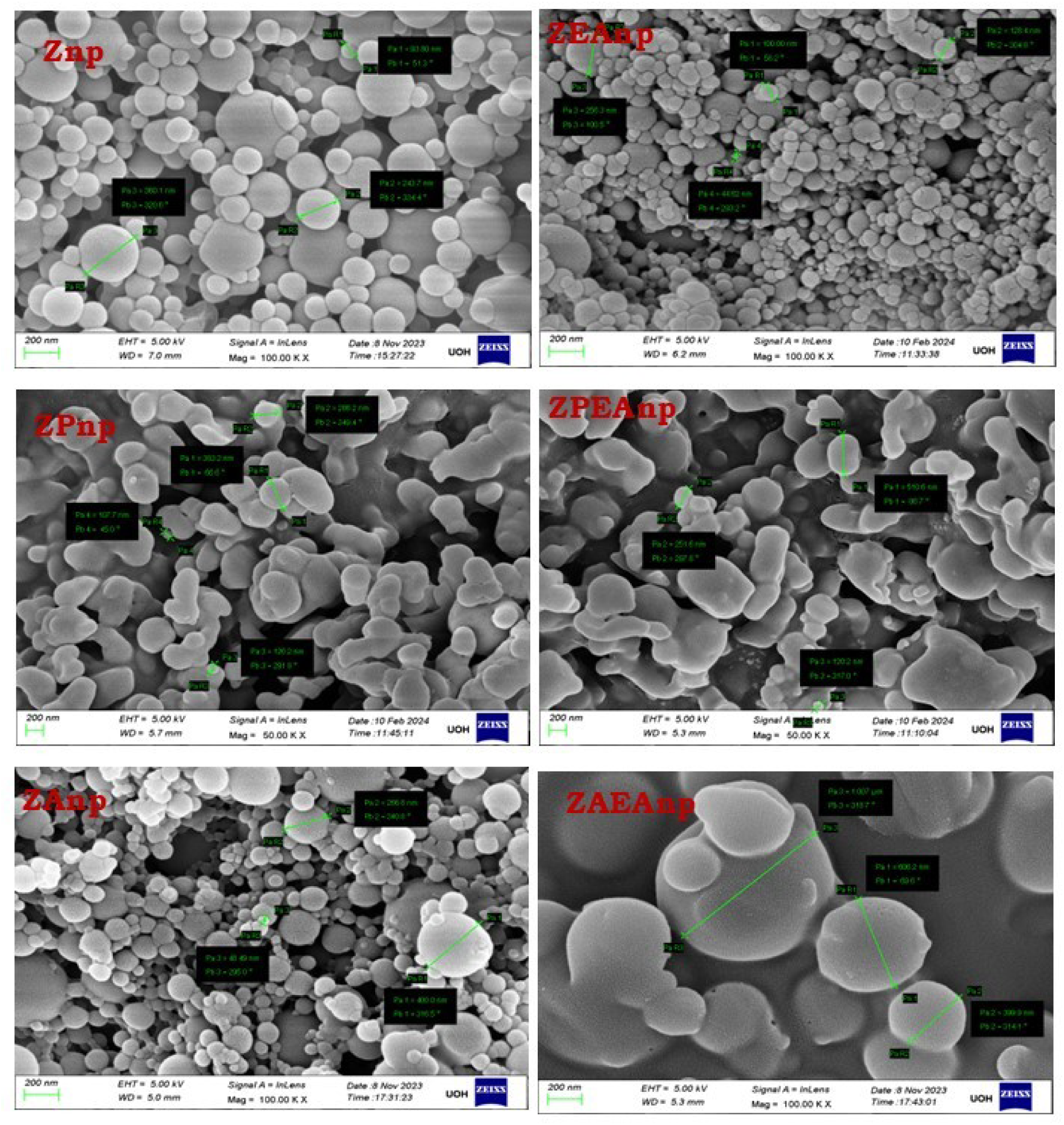
SEM image of the nanoparticles. Images a,c,e show Zein nanoparticles (**Znp**), Zein-Pectin nanoparticles (**ZPnp**), Zein-Alginate nanoparticles (**ZAnp**), and images b, d, and f show ethyl acetate extract (**EA**) encapsulated Zein nanoparticles (**ZEAnp**), Zein-Pectin nanoparticles (**ZPEAnp**), and Zein-Alginate nanoparticles (**ZAEAnp**)

**Table 2:** Size of nanoparticles.

Table No. 2:
| **SI No** | **Nanoparticles** | **Size range** |
| --- | --- | --- |
| **1** | Z-np | 93 – 360 nm |
| **2** | ZEAnp | 44 – 256 nm |
| **3** | ZP-np | 107 – 383 nm |
| **4** | ZPEAnp | 120 – 510 nm |
| **5** | ZA-np | 48 – 400 nm |
| **6** | ZAEA – np | 400 – 1007 nm |

### UV-Visible spectroscopic analysis

Figure 2a-c represents the overlay spectra of Znp and ZEAnp; ZPnp and ZPEAnp; ZAnp and ZAEAnp with their components. The result confirmed the incorporation of all the elements in the respective nanoparticles [35,36]. Figure 2a shows the overlay spectra of zein, ethyl acetate, zein nanoparticle, and extract-incorporated zein nanoparticle. The comparison of the spectrum of zein with Znp and ZEAnp confirms the presence of zein in both nanoparticles. The absorption maxima of zein at 280 nm may be due to the aromatic amino acid tyrosine’s *Ԉ-Ԉ\** transition [36], observed in both Znp and ZEAnp. The presence of ethyl acetate in ZEAnp was confirmed by studying the spectrum of ethyl acetate and ZEAnp. Two absorption bands with absorption maxima of 249 nm may be due to *Ԉ-Ԉ\** transition of the chromophore and 325 nm owing to the *n-Ԉ\** transition in the hetero atom were observed in the ethyl acetate fraction. Similar bands were observed in ZEAnp. The overlay spectra of zein, pectin, ethyl acetate, ZPnp, and ZPEAnp are indicated in Figure 2b. The absorption spectrum of ZPnp and ZPEAnp was taken in water and 80% ethanol, as the solubility of pectin was in water, and that of zein and ethyl acetate extract was in 80% ethanol. Comparing the spectrum of pectin with the ZPnp and ZPEAnp dissolved in water showed the absorption maxima of pectin at 204 nm, and were shifted to 223 nm; this may be due to the interactions with other components that are found common in ZPnp and ZPEAnp. The spectra of ZPnp and ZPEAnp in 80% ethanol when compared, observed that the absorption maxima of 280 nm. The ZPEAnp has exhibited a broad absorbance band ranging from 240 nm to 340 nm, possibly due to the presence of ethyl acetate extract, as it exhibits two absorption maxima at 240nm and 340 nm, respectively. Figure 2c shows the overlay spectra of zein, sodium alginate, ethyl acetate extract, Znp, and ZAEAnp. The spectra of ZAnp and ZAEAnp have been taken in both 80 % ethanol and water, as sodium alginate is soluble in water and zein in 80 % ethanol. The sodium alginate absorption maximum shifted from 197 nm to 251 nm, 220 nm ZAnp, and ZAEAnp, respectively, confirming the presence of sodium alginate in the nanoparticles. The overlay of spectra of zein with Znp indicates the presence of zein in Znp. The spectrum of ZAEAnp in 80% ethanol exhibited three prominent absorbance bands at 248, 283, and 316 nm, which may be due to the presence of ethyl acetate extract and zein, which show a broad absorbance band ranging from 253 - 353 nm and 257 - 333nm, respectively. The interactions between the components might cause the shift in the absorbance band of ZAEAnp from 253 nm to 248nm, with the emergence of a new band at 283nm, and another shift to 316nm from 353 and 333 nm, which was observed in Ethyl acetate extract, and zein [35,36].

**Figure 2:**
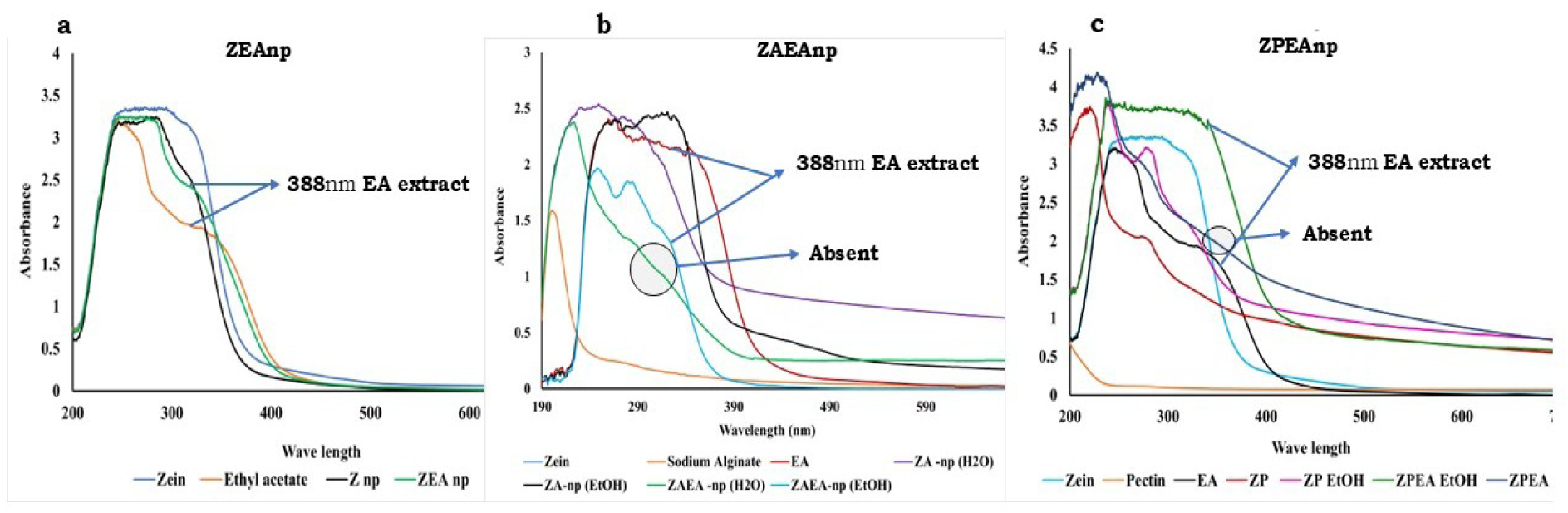
UV-Visible overlay spectra of different zein nanoparticles. Figure **a** shows the overlay spectra of Zein, Ethyl acetate, Znp, and ZEAnp; Figure **b** shows the overlay spectra of zein, ethyl acetate, sodium alginate, Zanp, and ZAEAnp in water and ethanol, and Figure **c** shows the overlay spectra of zein, ethyl acetate, pectin, ZPnp, and ZPEAnp in water and ethanol. It can be observed that the absorption band corresponding to EA is present in ZEAnp, ZAEAnp, and ZPEAnp dissolved in ethanol but is absent when the nanoparticles are dissolved in water.

### FT-IR spectroscopic analysis

Fourier Transform Infra-Red spectroscopic analysis validated the incorporation of zein, pectin, sodium alginate, and ethyl acetate extract components in respective nanoparticles, and an insight into the nature of their interactions. Figure 3-5 shows the FT-IR spectra of all the individual components and the nanoparticles with and without extract. Table 3 - 5 compares the peaks in the extract-encapsulated nanoparticles and their components. The peaks assigned for EA fraction 1363 cm^-1^ (-OH bending vibration of phenol), 1721 cm^-1^ (-CH bending of aromatic compounds), and 3332 cm^-1^ (-NH stretching vibrations of aliphatic primary amine) were found to be shifted to 1370 cm^-1^, 1737 cm^-1^, and 3315 cm^-1^ in ZEAnp [37–39]. This shift may be due to the interaction of phenolic -CH groups of aromatic compounds, and -NH groups of primary amines in the extract with zein, which confirms the incorporation of EA fraction in the nanoparticle [38]. In ZPEAnp, the respective peaks of EA were shifted to 1367 cm^-1^, 1733 cm^-1,^ and 3322 cm^-1,^ respectively, confirming the incorporation of the extract. The peak shifts of 1363 cm^-1^ to 1366 cm^-1^ and 1721 cm^-1^ to 1740 cm^-1^ confirmed the incorporation of the extract in ZAEAnp [40].

**Figure 3:**
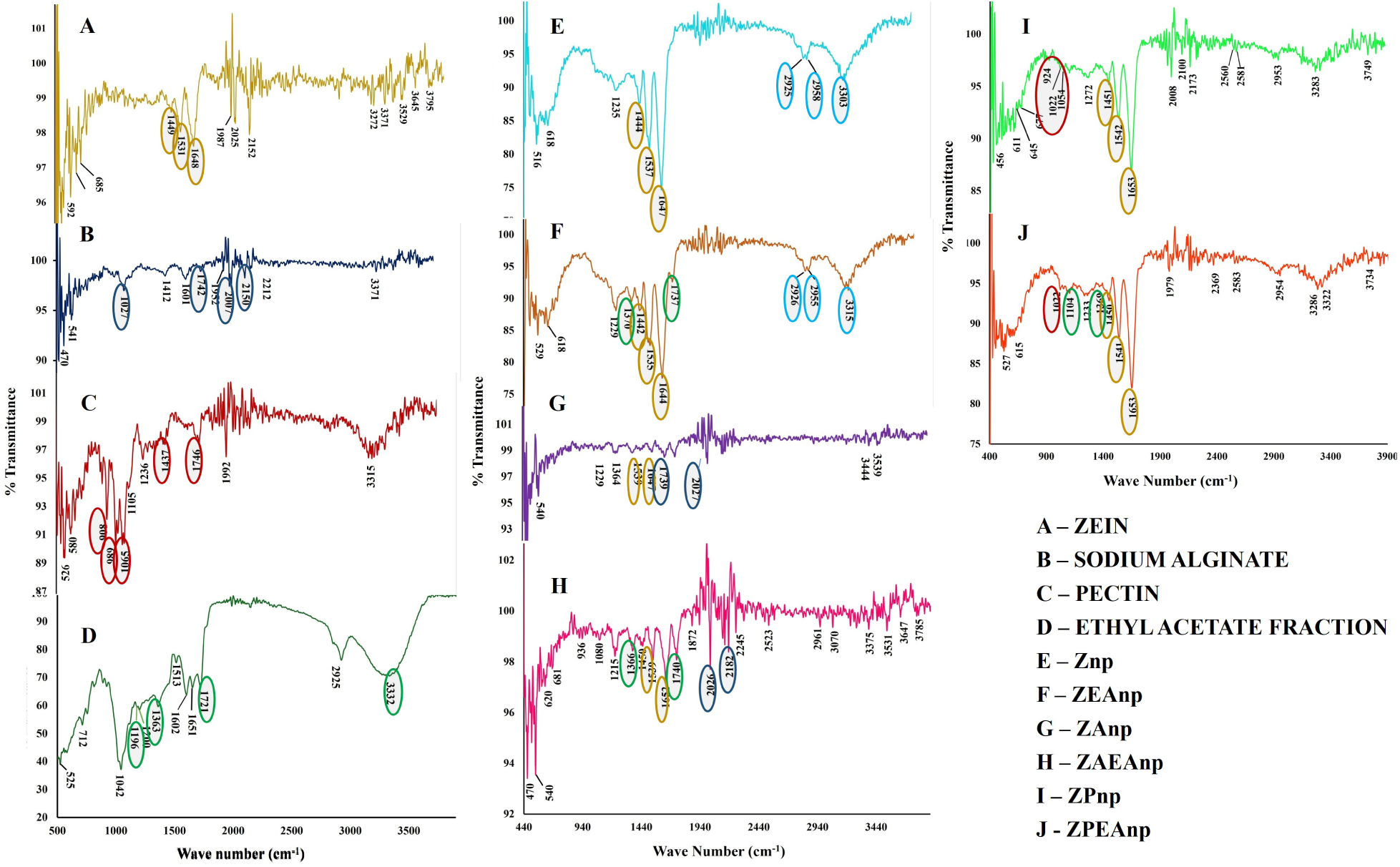
FT-IR Spectra of nanoparticles and their components. Figure A – D shows the FT-IR spectrum of the components of the nanoparticles: Zein, Sodium alginate, Pectin, and Ethyl acetate fraction, and E – J shows the spectra of Znp, ZEAnp, ZAnp, ZAEAnp, ZPnp, and ZPEAnp, respectively. The peaks and peak shift of zein (A) are denoted in yellow, which can be observed in Znp, ZEAnp, ZAnp, ZAEAnp, ZPnp, and ZPEAnp. Major peaks in Sodium alginate (B) are indicated in dark blue colour, and respective peaks and peak shifts in ZAnp (G) and ZAEAnp (H) are marked in the same colour. Peaks of pectin (C) and its shifts in ZPnp (I) and ZPEAnp (J) are indicated in red. The green mark indicates the characteristic peaks in EA (D) and its shift in ZEAnp (F), ZAEAnp (H), and ZPEAnp (J).

**Figure 4:**
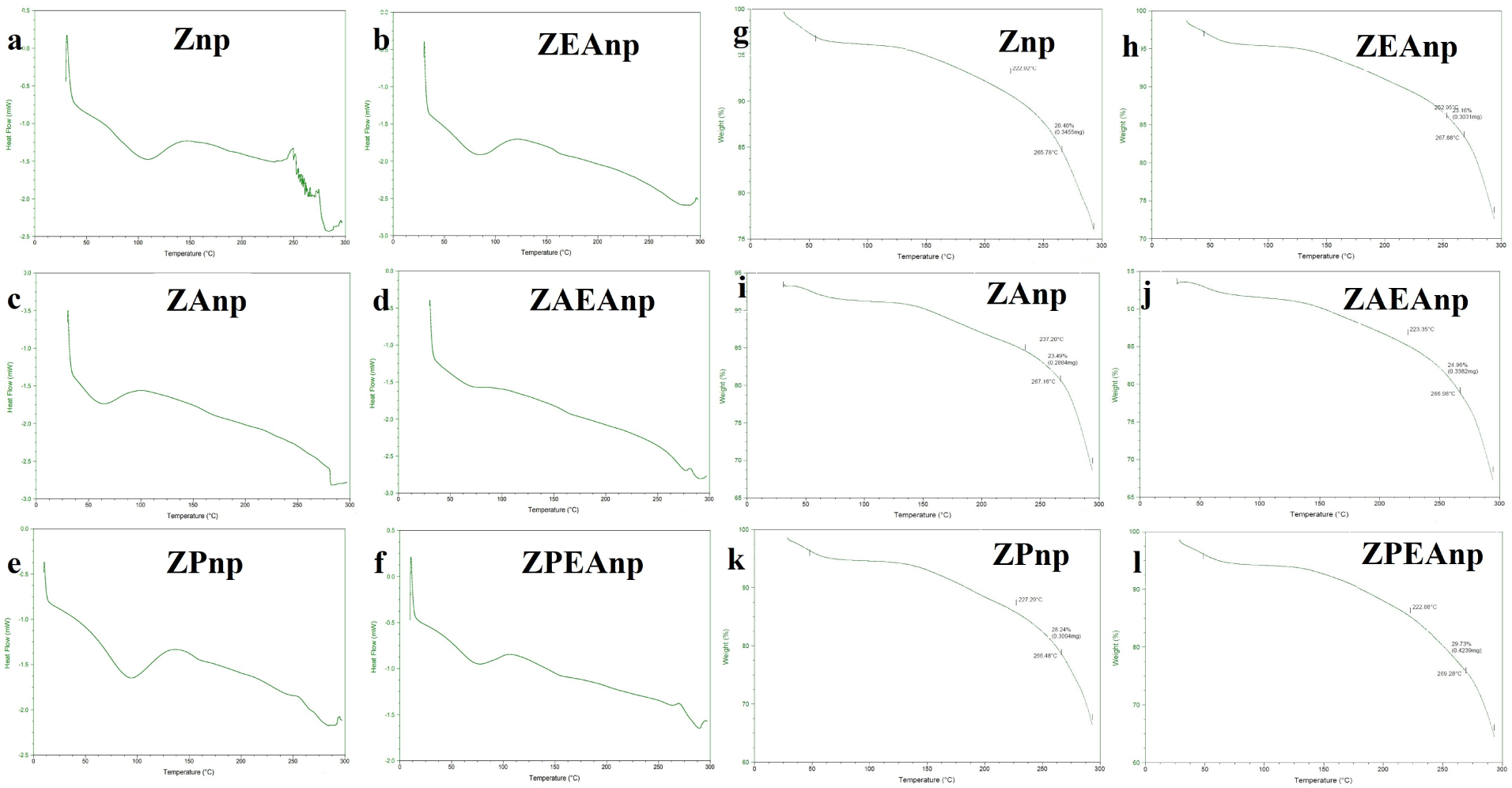
Thermal analysis of different zein nanoparticles. Figure (a - f) shows the DSC graphs, and (g – l) of Znp. ZEAnp, ZAnp, ZAEAnp, ZPnp, and ZPEAnp, respectively.

**Figure 5:**
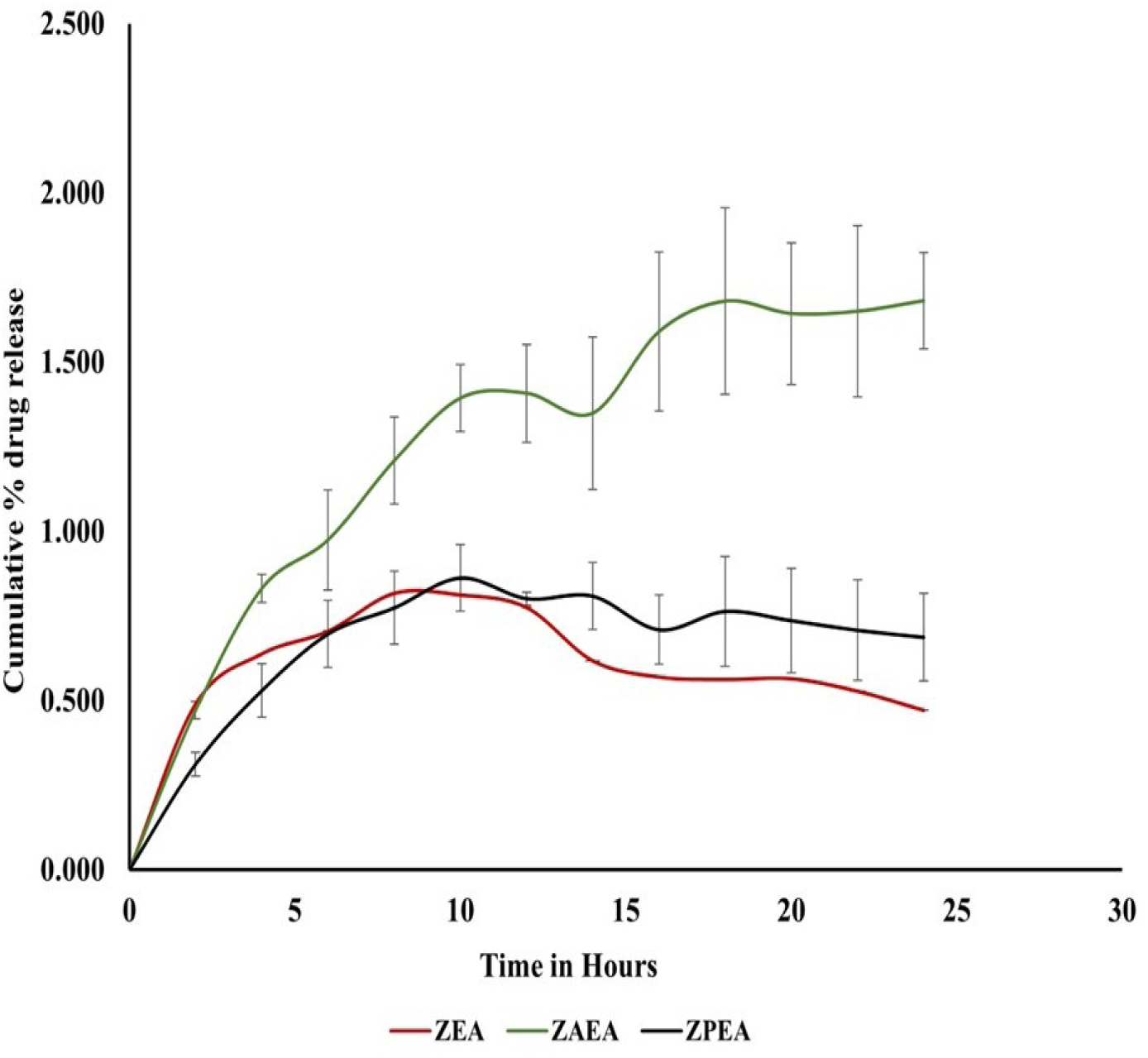
Cumulative percentage of drug release of different nanoparticles. ZAEAnp (green line) indicates a good amount of release compared to ZEAnp and ZPEAnp.

**Table 3:** Comparison of FT – IR peak values ZEAnp with its components.

Table No. 3:
| Ethyl Acetate Fraction<br>(cm<sup>-1</sup>) | Zein<br>(cm<sup>-1</sup>) | ZEAnp<br>(cm<sup>-1</sup>) | Inference |
| --- | --- | --- | --- |
|  |  | 1229 | -CO stretching vibration |
| 1363 |  | 1370 | -OH bending vibration of phenol |
|  | 1449 | 1442 | Alkane -CH<sub>3</sub> bending vibrations |
|  | 1531 | 1535 | -NH bending vibrations / -CH stretching vibration |
|  | 1648 | 1644 | -C=O- stretching vibrations of the peptide bond |
| 1721 |  | 1737 | -CO- group in dimerized aliphatic acids |
| 3332 |  | 3315 | Alkene -CH stretching vibrations |

**Table 4:** Comparison of FT – IR peak values ZAEAnp with its components.

Table No. 4:
| Ethyl acetate<br>(cm<sup>-1</sup>) | Zein<br>(cm<sup>-1</sup>) | Alginate<br>(cm<sup>-1</sup>) | ZAEAnp<br>(cm<sup>-1</sup>) | Inference |
| --- | --- | --- | --- | --- |
|  |  | 1027 | 1080 | C-O stretching vibrations in alcohols and phenols |
| 1363 |  |  | 1366 | O-H bending vibrations in alcohols & phenols |
|  | 1449 |  | 1450 | O-H bending vibrations in alcohols & phenols |
|  | 1531 |  | 1539 | -NH bending vibrations /-C-N stretching vibrations of amide groups |
|  | 1648 |  | 1652 | C=C stretching vibration of unconjugated linear alkenes |
| 1721 |  |  | 1740 | C=O stretching vibrations |
|  |  | 1742 | 1740 | C=O stretching vibrations in unsaturated aldehydes |

**Table 5:** Comparison of FT – IR peak values ZPEAnp with its components.

Table No. 5:
| **Ethyl acetate**<br>**(cm<sup>-1</sup>)** | **Zein**<br>**(cm<sup>-1</sup>)** | **Pectin**<br>**(cm<sup>-1</sup>)** | **ZPEAnp**<br>**(cm<sup>-1</sup>)** | **Inference** |
| --- | --- | --- | --- | --- |
|  |  | 1066 | 1021 | C-O stretching vibrations in alcohols & phenols |
|  |  |  | 1104 | C-O stretching in alcohols & phenols |
| 1363 |  |  | 1367 | -OH bending vibration of phenol |
|  |  | 1436 | 1450 | O-H bending vibrations in alcohols & phenols |
|  | 1531 |  | 1540 | -NH bending vibrations/C-N stretching vibrations of amide bonds |
|  | 1648 |  | 1651 | -C=O- stretching vibrations of the peptide bond |
| 1721 |  |  | 1733 | -C=O stretching vibrations in ketones |
|  |  | 1746 | 1733 | -C=O stretching vibrations in ketones |
| 3332 |  |  | 3322 | -N-H vibrations in amines |

### Thermal stability studies

The thermal stability of the nanoparticles was analyzed with thermogravimetric analysis (Figure 6) and differential scanning calorimetry (Figure 7). The thermogravimetric analysis observed that the first phase transfer occurs at 63.50 °C, 71.70 °C, and 60 °C for ZEAnp, ZAEAnp, and ZPEAnp, respectively. This may be due to the loss of water and alcohol, and it is found to be insignificant due to strong interactions with zein and alginate [41,42]. The degradation of ZEAnp and ZAEAnp starts at 259 °C, whereas the degradation of ZPEAnp starts at approximately 230 °C. The thermogravimetric analysis results indicated that all three nanoparticles exhibit good thermal stability.

**Figure 6:**
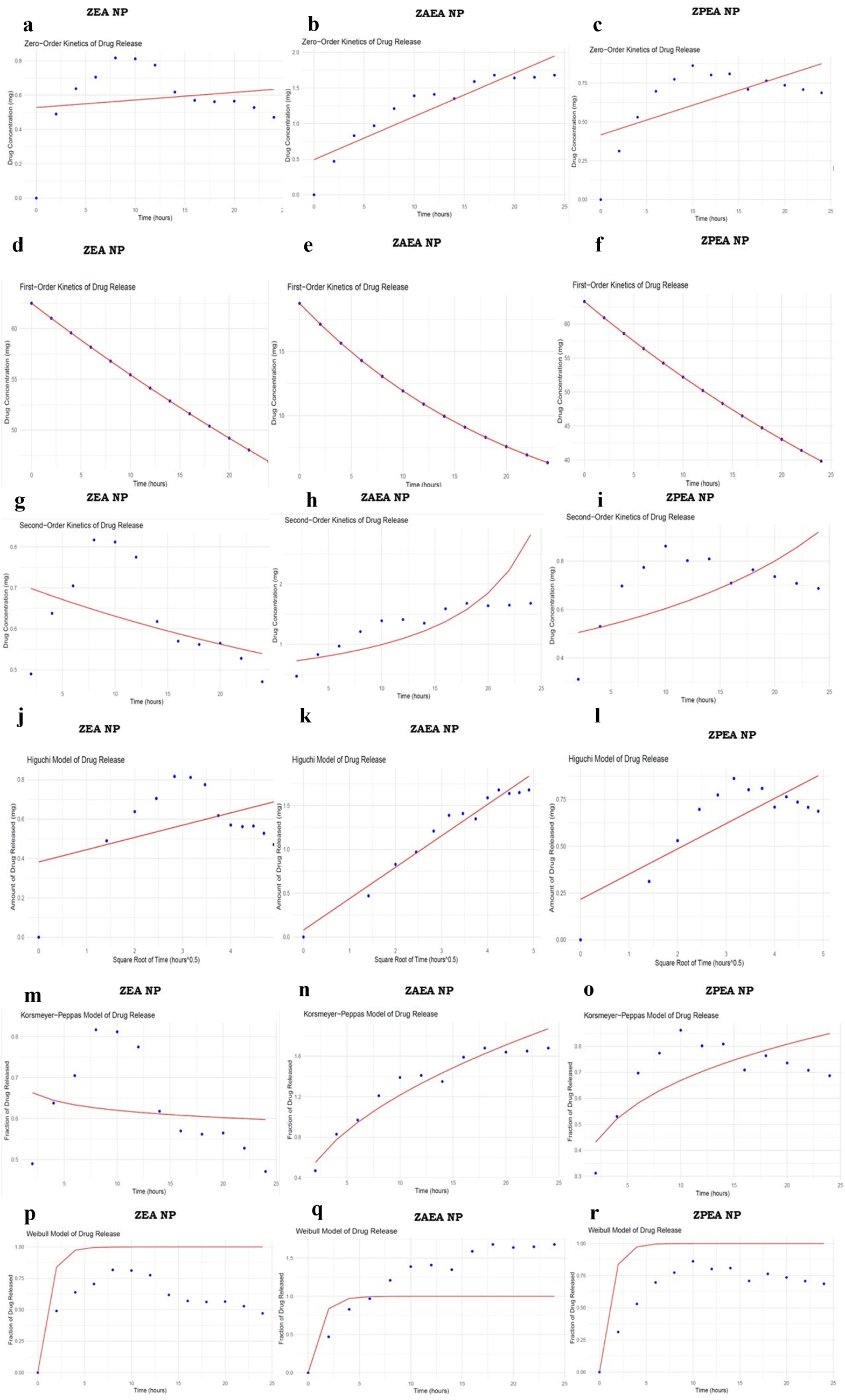
Drug release Kinetics of different nanoparticles. The figures (a-c) show Zero-order kinetics of drug release; (d–f) First-order kinetics of drug release; (g-i) Second-order kinetics of drug release; (j-l) Higuchi kinetics of drug release; (m-o) Korsmeyer Peppas model of drug release; (p-r) Weibull model of drug release of ZEAnp, ZAEAnp and ZPEAnp

The DSC curves of ZEAnp, ZAEAnp, and ZPEAnp indicate glass transition peaks at 88.5 °C, 69 °C, and 77 °C, respectively. For ZEAnp, the onset temperature of phase transition was 38.5 °C and endset was 115.4 °C with a temperature difference of 76.9 ^0^C. It can be assumed that the amorphous nature of ZEAnp undergoes a transition to a glassy form, but the crystallization begins only after 250 °C. The DSC curve of ZAEAnp indicated a glass transition at 69 °C with an onset temperature of 38.5 °C and an end set at 96.2 °C with a temperature difference of 57.7 ^0^C with less transition temperature. Hence, we can assume that only a minor change in structure occurs in ZAEAnp may be a transition from amorphous to rubbery form. Glass transition temperature at 77 °C was observed in ZPEAnp with an onset temperature at 38.5 °C and an end set at 108 °C. The temperature difference of 69.5 °C indicated the phase transition from its amorphous nature to a rubbery or glassy form. The thermal stability studies indicated that all three nanoparticles exhibit considerable thermal stability and are stable at room temperature.

### *In-vitro* drug release

The drug release pattern of the three nanoparticles ZEAnp, ZAEAnp, and ZPEAnp exhibited a sustained release. However, the cumulative % of drug release in ZEAnp and ZPEAnp was found to be very low, and at the 12^th^ hour, the drug release from ZEAnp started to decline, whereas in ZPEAnp, till the 10^th^ hour, there was a steady rise in the release and almost attained equilibrium after the 18^th^ hour. The ZAEAnp exhibited an exceptional pattern of drug release with a slow and steady rise in drug release. From the drug release plot, it is evident that ZAEAnp exhibits 1.5% of drug release at the 20^th^ hour, and a steady state is attained after that. These results indicate a drug release pattern for a prolonged period. To understand the drug release patterns, the drug release data were fitted into different mathematical models, such as zero, first, second, Higuchi, Korsmeyer Peppas, and Weibull models. The release pattern was best fitted in first-order, Higuchi, Korsmeyer, Peppas, and Weibull models. Even though all the nanoparticles obey first-order kinetics, which indicates Fickian diffusion-based drug release, the Higuchi and Korsmeyer Peppas models show that ZAEAnp exhibits better release properties than ZEA and ZPEAnp. The Higuchi dissolution constant ‘k’ was found to be high for ZAEAnp. The higher the ‘k’ value, the faster the release. Accordingly, the drug release rate was in ZAEA np>ZPEA np>ZEA np. The release exponent ‘n’ was calculated in the Korsmeyer Peppas model. If the ‘n’ value is less than 0.5, it indicates Fickian diffusion, where the drug moves from a region of its higher concentration to its lower concentration. The ‘n’ value of ZEAnp, ZAEAnp, and ZPEAnp was found to be -0.042, 0.49, and 0.27, respectively. From this result, it was confirmed that the ZEAnp does not exhibit good drug release and cannot be used as a drug carrier. ZAEAnp and ZPEAnp had ‘n’ values less than 0.5 and hence can be used as drug carriers. However, ZAEAnp shows better drug release properties when compared to ZEAnp and ZPEAnp as its ‘n’ value is almost near to 0.5 and has the highest ‘k’ value from Higuchi kinetics. From the results, it can be concluded that incorporating sodium alginate in zein nanoparticles enhances its stability, encapsulation efficiency, and drug release [11,30,31].

### Conclusion

This study encompasses the development of different zein nanoparticles with the encapsulation of herbal extract in a green synthetic approach, namely ZEAnp, ZAEAnp, and ZPEAnp. The morphological and structural studies indicate the formation of spherical nanoparticles and the incorporation of all the components in the nanoparticles. The thermal studies envisage that the nanoparticles are stable at room temperature. It was also evident from the DSC curve that the encapsulation of ZAnp with EA extract to form ZAEAnp improves the thermal stability of the same. Even though the ZEAnp exhibited good morphological and thermal characteristics, the encapsulation efficiency and drug release properties were found to be low. Our research confirms that among the two different polysaccharides, sodium alginate and pectin used for modifying the zein nanoparticle (Znp), the incorporation of the anionic polysaccharide sodium alginate improves the encapsulation efficiency and drug-release properties of zein nanoparticles.

## Author contributions

DAT: Performed the experiments analyze the results and wrote the manuscript; VN: Performed experiments helped in analyzing results. AM: Performed experiments and helped in analyzing results SRK: Conceptualized, designed the experiments, interpreted the data, supervised the entire work, and edited the manuscript.

## Conflict of Interest

None

## Acknowledgement

We acknowledge the financial support of Anusandhan National Research Foundation (ANRF) Govt. of India (ANRF/IRG/2024/001352/LS), Kerala State Council for Science, Technology and Environment (KSCSTE∕1095∕2020-PDF) and acknowledge the DST-FIST, infrastructural facilities of the department. DAT: thankful to Kerala State Council for Science, Technology and Environment for Post Doctoral Fellowship (KSCSTE∕1095∕2020-PDF).

## Notes

### Competing Interest Statement

The authors have declared no competing interest.

